# Correspondence between functional scores from deep mutational scans and predicted effects on protein stability

**DOI:** 10.1101/2023.02.03.527007

**Authors:** Lukas Gerasimavicius, Benjamin J Livesey, Joseph A. Marsh

**Affiliations:** MRC Human Genetics Unit, Institute of Genetics & Cancer, University of Edinburgh, Edinburgh, UK

## Abstract

Many methodologically diverse computational methods have been applied to the growing challenge of predicting and interpreting the effects of protein variants. As many pathogenic mutations have a perturbing effect on protein stability or intermolecular interactions, one highly interpretable approach is to use protein structural information to model the physical impacts of variants and predict their likely effects on protein stability and interactions. Previous efforts have assessed the accuracy of stability predictors in reproducing thermodynamically accurate values and evaluated their ability to distinguish between known pathogenic and benign mutations. Here, we take an alternate approach, and explore how well stability predictor scores correlate with functional impacts derived from deep mutational scanning (DMS) experiments. In this work, we compare the predictions of 9 protein stability-based tools against mutant protein fitness values from 45 independent DMS datasets, covering 161,441 unique single amino acid variants. We find that FoldX and Rosetta show the strongest correlations with DMS-based functional scores, similar to their previous top performance in distinguishing between pathogenic and benign variants. For both methods, performance is considerably improved when considering intermolecular interactions from protein complex structures, when available. Finally, we also highlight that predicted stability effects show consistently higher correlations with certain DMS experimental phenotypes, particularly those based upon protein abundance, and, in certain cases, can be competitive with other sequence-based variant effect prediction methodologies for predicting functional scores from DMS experiments.

## Introduction

Recent decades have seen massive breakthroughs in optimizing genomic sequencing for large-scale operations, revealing the high prevalence of genetic variation in human populations^1,2^. Many genetic variants are from missense mutations, which cause a change in the identity of single amino acid residues at the protein level^3^. However, the precise phenotypic consequences of most variants remain uncertain as mutants are seldom functionally characterized in clinical settings, while alternative approaches to conclusively classify variants, such as genetic testing and pedigree studies, require many cases^4^.

Multiplex assays of variant effects (MAVEs) have emerged as methodologies with the potential to measure the effects of large numbers of genetic variants in parallel within a single experiment^5,6^. MAVEs produce interpretable variant-function maps by associating each variant to a quantitative assay measurement for a select phenotype. MAVEs that involve protein-based assays of amino acid variants are often referred to as deep mutational scanning (DMS) experiments. DMS has been widely adopted to explore effects of amino acid variation using a variety of different experimental phenotypes, such as protein abundance, activity or general cellular fitness^5^. However, while the number of proteins that have been characterized through DMS grows constantly^7,8^, and use of these methodologies and coordination between groups is increasing through the Atlas of Variant Effects Alliance^9^, as of now DMS is not up to the challenge of evaluating all possible substitutions in the entire proteome, both due to costs and the inherent limitations of assaying specific phenotypes.

As an alternative or complement to experimental approaches for characterizing variants, considerable efforts have been put into developing generalizable computational models for predicting the effects of protein variants. A large number of variant effect predictors (VEPs) exist that leverage different properties, including evolutionary sequence conservation, phylogenetic relationships and physicochemical properties, to evaluate the likelihood of a variant being damaging^10^. However, the scores output by these VEPs seldom provide an interpretable context to the underlying disease mechanism. An alternative approach is presented by structure-based protein stability predictors, which can evaluate the change in Gibbs free energy of folding (ΔΔG) or intermolecular interaction upon mutation^11^. Stability predictors are frequently used in the fields of protein engineering and even clinical genetics, despite not being trained for disease identification, because they can distinguish between stabilizing and destabilizing energetic effects and provide clues as to possible pathogenic mechanisms^12^.

The methodological approaches to predicting stability impacts of mutations are diverse. FoldX^13^ and Rosetta^14^ use empirical physics-based potentials with additional statistical terms based on observations from bimolecular structures. ENCoM is a unique method that takes into account how mutations can impact protein dynamics and stability through normal mode analysis^15^. A combination of evolutionary and structural information has been employed in the untrained DDGun3D predictor^16^. Some recent predictors, like mCSM, have been derived through machine learning using various features^17^. Given such heterogeneity in approaches, numerous studies have been carried out to benchmark the performance of these predictors in reproducing realistic ΔΔG values that agree with experimental thermostability data^11,18–23^. Furthermore, there have been attempts to explore and address biases, such as data circularity, overprediction of destabilizing variants and lack of prediction symmetry^16,24–27^. However, as stability predictors are now routinely used for protein engineering and disease identification purposes^28–35^, it is crucial to know how well ΔΔG serves as a proxy score for pathogenicity, and thus how prevalent are destabilizing loss-of-function mechanisms in the pool of all possible mutations^36–38^. We have previously assessed the performance of ΔΔG values from stability predictors in distinguishing between pathogenic and putatively benign missense variants in a classification task^12^. Phenotypic assays now offer further opportunity to more quantitatively interrogate the extent to which predicted ΔΔG agrees with assayed fitness or activity of protein variants.

DMS datasets currently provide the most extensive experimentally derived representation of the functional variant effect landscape, and they have been very successfully utilized in recent VEP benchmarking studies^39,40^. It was further shown that DMS assay values themselves can, in some cases, be better at distinguishing pathogenic from benign variants than current computational approaches. However, a number of caveats should be understood when using DMS scores to evaluate how damaging effects are represented through predicted changes in stability. Realistically, we can expect that assays that evaluate phenotypes such as protein abundance or complex formation, should show the best agreement with stability predictions, as they are well suited to detect destabilizing loss-of-function molecular mechanisms. Other types of assays, such as general competitive growth experiments, are potentially sensitive to non-destabilizing but damaging mutations, such as those associated with gain-of-function or dominant-negative effects. We have previously shown that damaging mutations, which manifest through such non-loss-of-function mechanisms, tend to be mild at a protein structural level, and not well identified through prediction of stability effects^38^. Thus, some specific types of assays, which do not measure stability directly, may show very heterogenous agreement with stability predictors. However, we believe DMS values provide an unbiased, independent way of comparing predictors, and at the same time allow us to explore how well destabilizing loss-of-function mechanisms can be identified through specific experimental phenotypes.

In this study, using a large number of DMS datasets as a benchmark, we quantified the capability of structure-based protein stability predictors to accurately rank the functional impacts of variants. We demonstrate that FoldX and Rosetta predictions derived on protein complex structures significantly outperform other tools in assessing the functional impact of mutations. We also show how evaluating full biological assemblies improves our ability to relate predictions to functional phenotypes involving protein or DNA binding. Finally, we explore how certain types of DMS phenotypes, specifically ones related to protein abundance, correlate the best with variant stability predictions due to their closer association with destabilizing loss-of-function mechanisms.

## Results

### Considerations of stability predictor benchmarking with DMS datasets

For this study, we gathered 45 different DMS datasets for 35 unique protein targets. A majority of the datasets we used were collected and outlined previously in the comprehensive VEP benchmarks from Livesey & Marsh^39,40^. This mostly included assays of human proteins, but also experiments on yeast, bacterial and viral proteins. In addition, new datasets have been subsequently released and published in the MaveDB^7^, which were combined with previous data to form our benchmarking dataset. The full list of genes and DMS datasets used in this study is available in **Supplementary Table 1.** For the additional DMS experiments that contained fitness values from assays under multiple conditions, we selected the option most representative of native-like conditions, for instance grown under no additional treatments than were required by the phenotypic assay or experimental setup.

The DMS datasets used are heterogenous in terms of the phenotypes that were assayed for. The most frequent category involves human gene complementation of native genes in yeast, with the fitness scores being derived from competitive growth of variants in deep mutational scanning experiments. VAMP-seq is another methodology, which interrogates the abundance of GFP-fused variant proteins and thus also their stability^41^. Other methods include toxicity assays, assessing changes to binding strength through two-hybrid or phage display assays, or activity assays tailored to specific targets, for instance patch-clamp assessment of cell currents for KCNQ4 potassium channel variants^42^. In this work, for cases where a single gene has multiple associated DMS datasets, we identify the datasets alphabetically, surrounded by parentheses, *e.g*. BRCA1(a) and BRCA1(b) represent two independent DMS experiments performed by different groups using different functional assays.

Different DMS datasets for the same genes can show variability in variant impact values due to different experimental conditions or even the phenotypes being assayed, however, they generally show moderate to high Spearman’s correlations, averaging at around 0.66, suggesting that they represent a sufficiently robust approach to benchmark variant effect prediction performance^39^. Mutations that excessively perturb the stability of a protein and lead to its degradation ought to be reflected by most assay types. However, we can also imagine that many DMS assays would be sensitive to mutations that affect the protein in some way other than loss of function due to intra- or inter-molecular destabilization. Thus, the degree of agreement between ΔΔG values and DMS scores should also tell us something of the pervasiveness of destabilizing loss-of-function mechanisms for a given gene or tested phenotype.

Most of the stability predictors we use here depend on structural inputs. We therefore derived a structural variant map, linking each variant in every DMS dataset to a residue within a protein structure, using the same strategy as described recently^38^. AlphaFold2 models were used for proteins or select residues in the infrequent cases where they were not covered by experimental structures. It has been recently demonstrated that accurate stability predictions can also be delivered using modelled protein structures^43,44^.

We tested 9 stability predictors, 7 of which we have previously also explored for their ability to distinguish between pathogenic and putatively benign human variants^12^. On top of FoldX, Rosetta, INPS3D, PoPMuSiC, mCSM, ENCoM and DynaMut2, we have included DDGun3D, an ‘untrained’ stability prediction method, as well as the recently released RaSP which offers rapid evaluation of variants based upon sequence alone through a neural network model^13–17,44–47^. While most methods only offer functionality of evaluating stability perturbing effects of mutations on monomeric structures, FoldX, ENCoM and Rosetta were also evaluated in terms of full protein complex structures, if they were available, as this functionality is easily accessible in these predictors. Finally, we not only explored the agreement of ΔΔG values, which range from stabilizing to destabilizing, but also absolute change in stability, | ΔΔG |, as a metric of general energetic perturbation of the structure. We have previously shown that considering strongly stabilizing variants as deleterious through use of | ΔΔG | improves the disease identification performance of most stability predictors, likely due to stability predictors sometimes mispredicting the sign of the ΔΔG, and stability-increasing mutations occasionally being pathogenic^12^.

Before benchmarking the predictors on DMS measurements, we first compared the agreement between all predictors for the full dataset of explored variants to get a sense of the heterogeneity between different methodological approaches. We calculated pairwise Spearman’s rho values across all evaluated variants between each predictor pair, using both ΔΔG and |ΔΔG| metrics **(Figure 1**). Without taking the high correlation between monomeric *vs* complex predictions for the same predictor into account, or DynaMut2 and its methodological component mCSM, we see that, overall, most of the predictors also show fairly good agreement with each other, with the average absolute Spearman’s rho value for ΔΔG at 0.49, rising to 0.62 if we exclude ENCoM.

**Figure 1.**
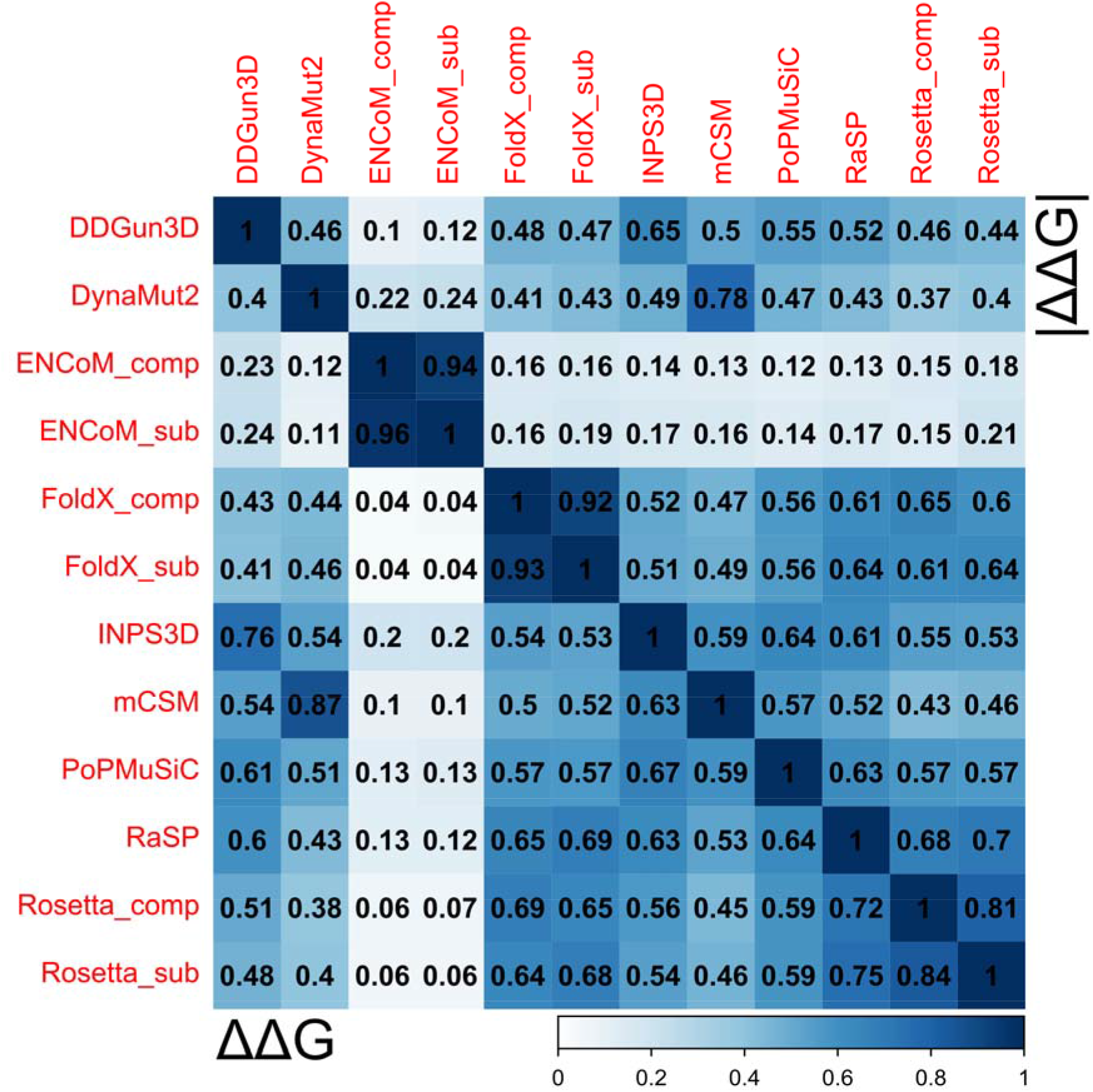
Most protein stability prediction methods display moderate agreement with each other. The coloured scale bar represents the absolute magnitude of Spearman’s rho values from pairwise comparisons calculated for complete observations, calculated across variants pooled from all the genes. The lower matrix triangle shows Spearman’s correlation calculated using the raw Gibbs free energy values, taking into account both destabilizing and stabilizing effects, while the upper triangle contains rho values derived using only the absolute magnitudes. All predictor ΔΔG values were adjusted to match for stabilizing and destabilizing effect directions.

ENCoM predictions stand out as the most unrelated to any other predictor, with the highest correlation for ΔΔG being with DDGun3D at 0.24. Rosetta predictions appear to be the most divergent in terms of comparing monomeric *vs* complex values for the same mutations, likely as a result of the energetic minimization procedure of structures, treating complexes as a single chain. As in our previous work, using absolute stability values did not prove beneficial to increase the agreement between most predictors, with the exception for ENCoM correlations with FoldX and Rosetta. We have also previously demonstrated that ENCoM benefits the most from use of absolute values for classification tasks, as it tends to overpredict stabilizing effects. The highest correlations of 0.75 were observed between DDGun3D and INPS3D, as well as between RaSP and Rosetta. Indeed, one would expect RaSP and Rosetta to show a high degree of correlation, as RaSP has been trained not on experimental ΔΔG values, but on Rosetta predictions^44^. The high correlation between DDGun3D and INPS3D is likely due to the fact that both methods are essentially extensions of their underlying sequence and alignment-based models, and therefore both directly utilize information about evolutionary sequence conservation, in contrast to all other predictors.

### Correspondence between stability predictions and DMS values

For each stability predictor and DMS dataset pair, we calculated Spearman’s correlations using the shared subset of observations **(Figure 2).** Both the predictor and DMS score directions were adjusted for consistency *i.e*. higher ΔΔG values indicate increased destabilization and higher DMS scores indicate more damaging effects). While we observed that most predictors rank DMS dataset variants with moderate consistency between each other, the overall correlations tend to be fairly low, with the average rho values throughout the benchmark at 0.26 and 0.28 for ΔΔG and | ΔΔG |, respectively. This is in-line with the recent observations from Høie *et al*. showing an average Spearman’s correlation value of 0.25 between Rosetta ΔΔG and a number of DMS datasets^48^. Overall, even direct experimental thermostability values and computational predictions are often shown to only correlate on the order of ~0.5, with some works showing that the practical upper bound for such comparisons can only reach ~0.8 due to experimental data quality^49^.

**Figure 2.**
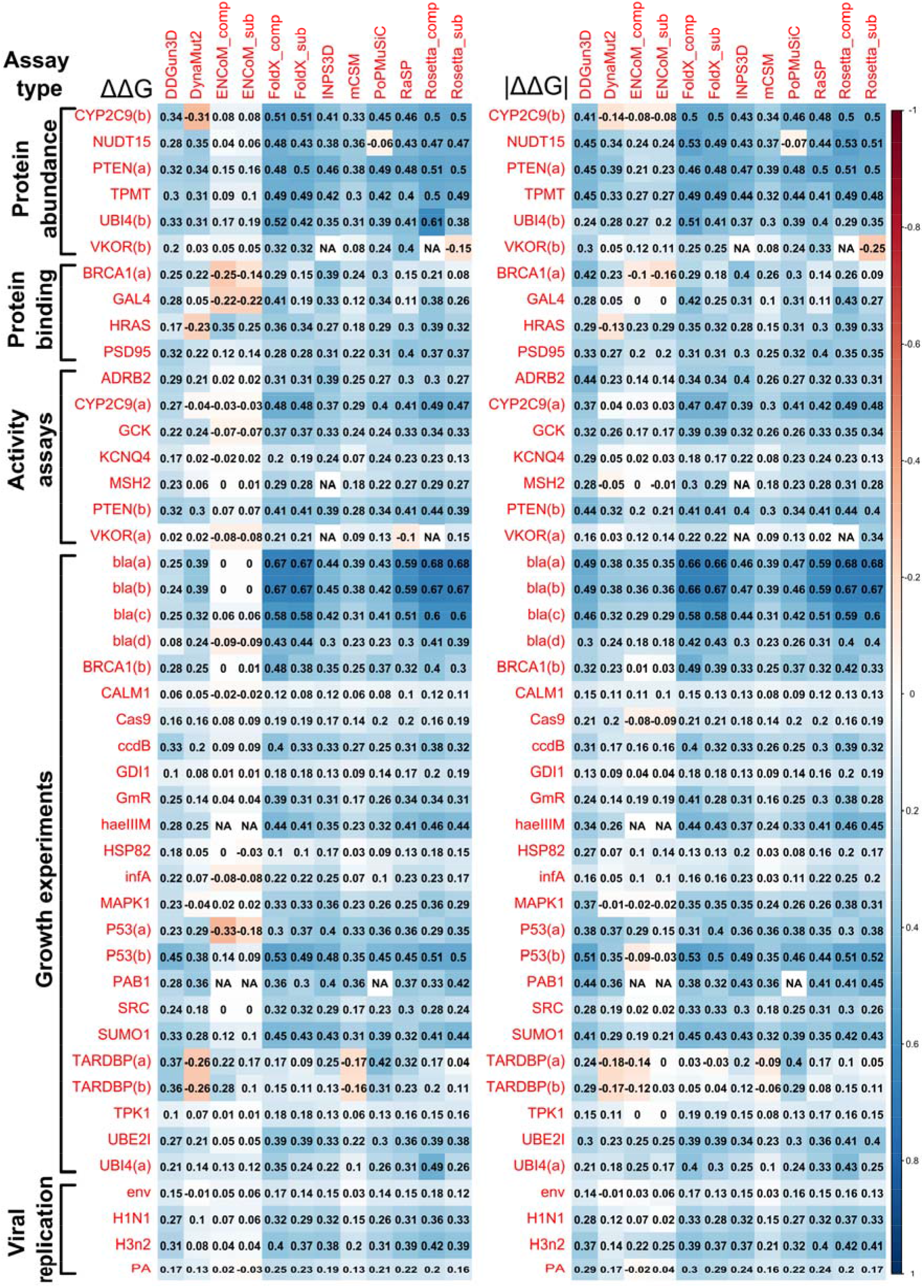
Stability predictor scores can correlate highly with scores from DMS experiments, but performance is highly heterogenous. The coloured scale bar represents the absolute magnitude of Spearman’s rho values from pairwise comparisons calculated for complete observations. All predictor ΔΔG values were adjusted to match for stabilizing and destabilizing effect directions.

However, other works have shown individual correlations between stability predictions and DMS functional scores can reach up to 0.57 for targets like NUDT15^50^. In our analysis, Rosetta and FoldX stability predictions for the *E. coli* beta lactamase (bla) antibiotic resistance DMS datasets bla(a) and bla(b) produced the highest observed correlations in this benchmark, ranging between 0.66-0.68, while the best correlation for bla(d), a different antibiotic resistance experiment, was only 0.44. Rosetta complex ΔΔG values also appear to correlate well (0.61) with dataset UBI4(b), based on a FACS assay that relates well to stability. The next best correlating datasets for other genes had Spearman’s rho values of up to 0.53, across both ΔΔG and | ΔΔG |.

We have previously shown that dominant-negative and gain-of-function disease variants tend to be structurally milder and demonstrate weak stability perturbation^38^. As such, they are harder to distinguish from benign variants through stability prediction. This is likely to contribute to the low correlations observed for CALM1, TARDBP and SRC datasets, which are known to also be associated with non-loss-of-function disease mechanisms, such as the dominant-negative effect in CALM genes^51–53^, and gain-of-function mutations in TARDBP^54,55^ and SRC^56,57^.

ENCoM, the method with weakest correlations overall, leverages a unique prediction approach based on normal mode analysis, which is purported to take into account variant effects on protein dynamics. However, it also benefits the most out of all predictors from the use of absolute stability values as a metric, increasing Spearman’s rho by up to 0.36, compared to correlations achieved using raw stability values. This approach shifts the perspective towards evaluating mild vs perturbing effects, instead of stabilizing vs destabilizing, suggesting a likely tendency to overpredict variants as stabilizing. This also appears to affect DDGun3D to a lesser extent.

Importantly, we also explored how taking into account intermolecular interactions, by using complex structures, impacts variant stability prediction, and in turn the evaluation of functional effects. We compared results from FoldX, Rosetta and ENCoM, which were the only methods that allowed easy use of biomolecular complex structures in prediction. This led to considerable improvements in the agreement with DMS measurements in certain cases. For example, both FoldX and Rosetta showed marked increases in correlation with GAL4 DMS values when the full DNA-bound structure was utilized, as opposed to just the monomeric subunit in isolation. This is illustrated in **Figure 3,** where the structure of the GAL4 complex is shown coloured based on DMS values, showing that hotspots of highest functional variant impact appear not only at the protein dimer interface, but also at interaction sites with DNA. Such mutation effects at interfaces would be missed by stability predictions that do not take intermolecular interactions into consideration.

**Figure 3.**
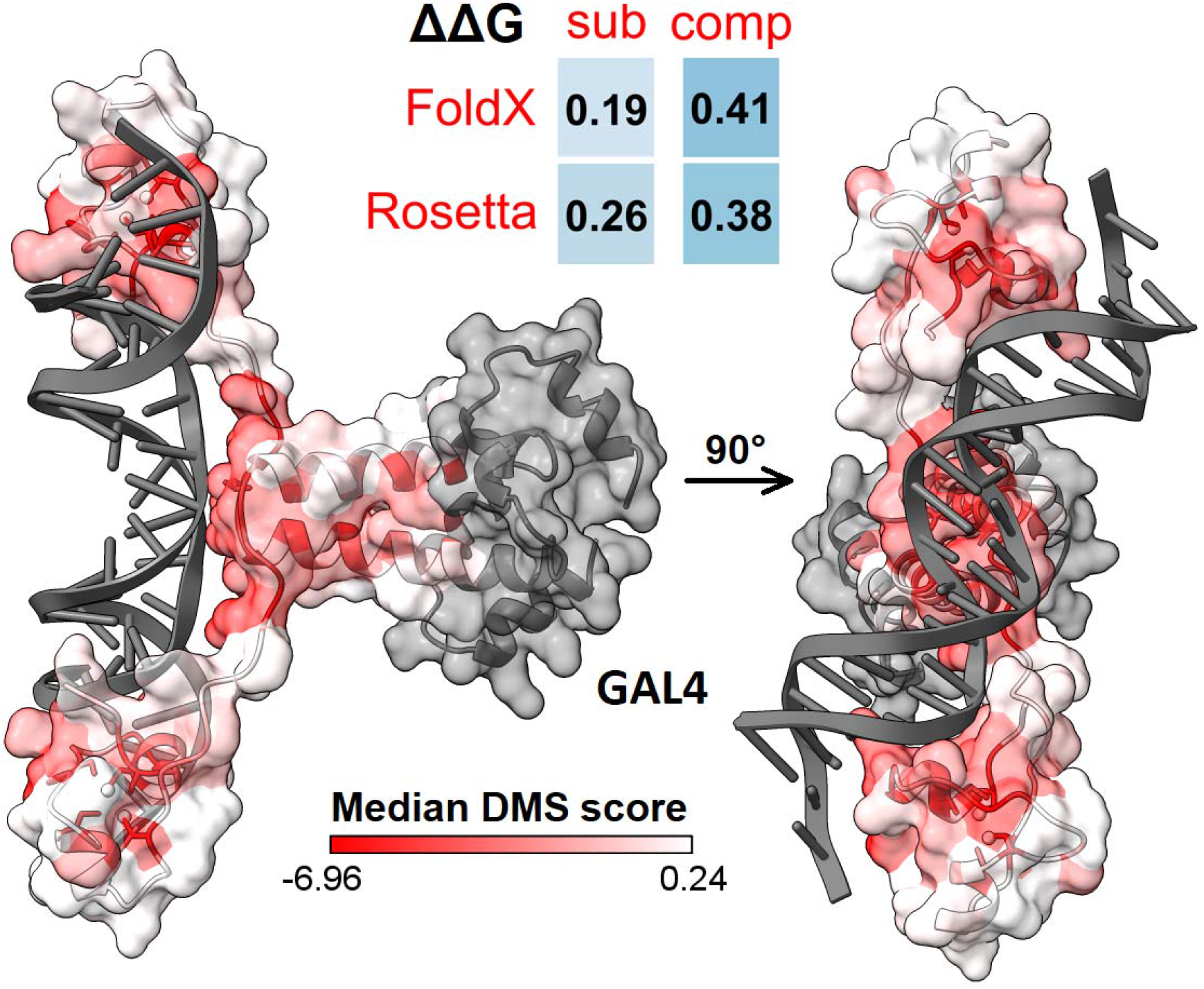
Stability predictors that can account for intermolecular interactions between proteins and other molecules, such as DNA, allow to explain a larger extent of functional variant effects through the scope of stability. Values in squares represent Spearman’s correlations for GAL4 variants between predicted Gibbs free energy changes and DMS scores, in the case of using just one subunit structure, or evaluating stability on the entire complex assembly. The DMS dataset for GAL4 contains variant enrichment scores from a yeast two-hybrid experiment. Protein structures are coloured based on the per-position median variant score, with lower values indicating more sensitive positions, and grey indicating missing values. GAL4 variants used for the analyses were mapped to PDB structure 3COQ, containing a GAL4 dimer and a DNA double-strand.

### Relative ranking of stability predictor performance in assessing variant functional impacts

To compare the relative performance of our tested predictors, we used a scoring and ranking scheme based on pairwise predictor correlation comparisons on each DMS dataset, as recently introduced^40^. Each predictor would be given a point for demonstrating a higher Spearman’s rho value for a DMS set against another prediction tool, or both would get half a point for a tie. The advantage of this ranking strategy is that it accounts for the fact that not all methods successfully made predictions for all mutations in all proteins, as the pairwise comparisons are performed only on the shared variant subset that both predictors were successfully able to evaluate. In the end each, predictor’s score was scaled by the total number of successful comparisons it had been involved in, resulting in a relative performance ranking, for both ΔΔG and | ΔΔG | **(Figure 4).**

**Figure 4.**
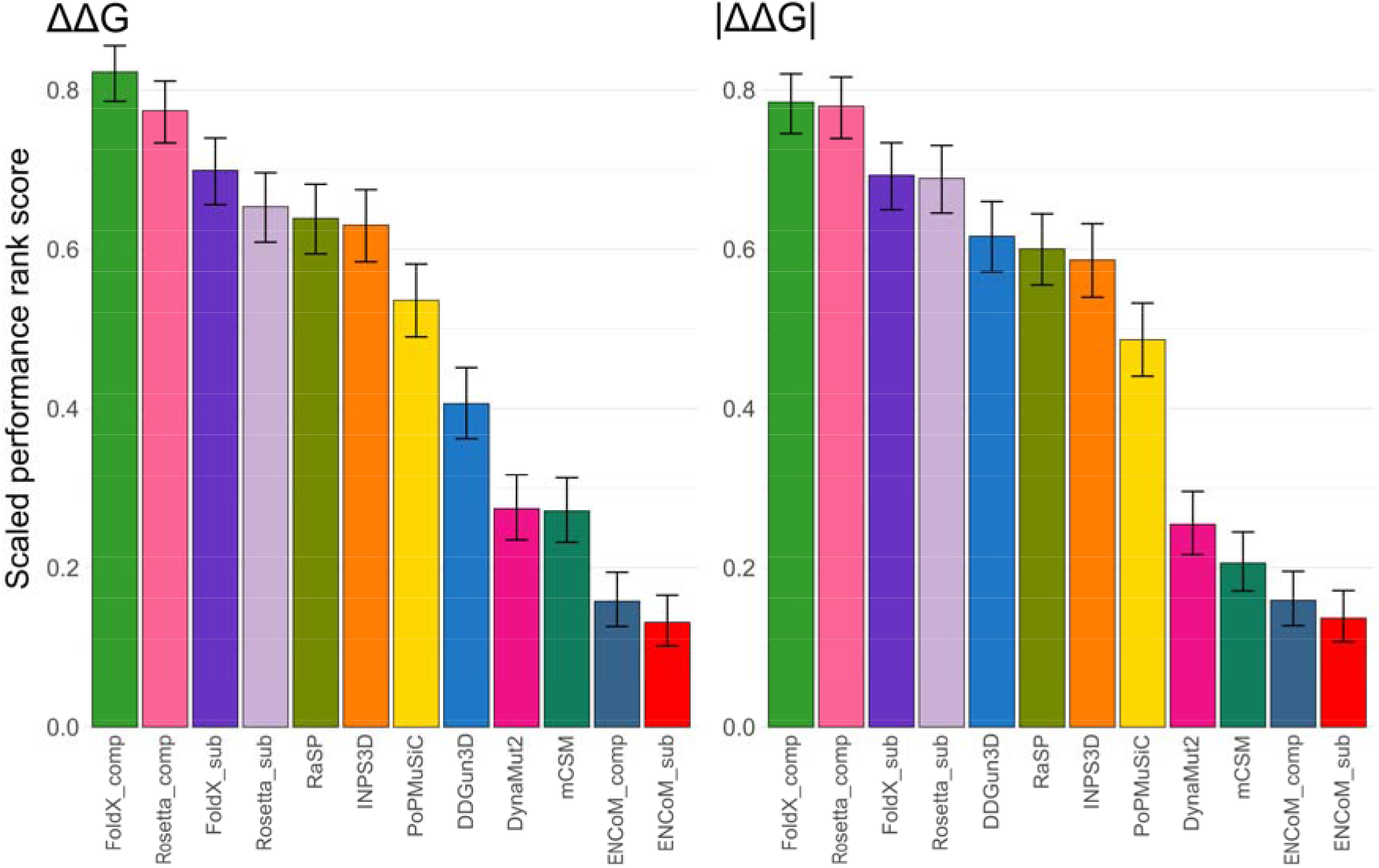
Complex-based FoldX and Rosetta stability predictions are significantly more accurate at ranking variant effects in relation to DMS scores than other stability evaluation tools. Computational protein stability predictor rankings were derived based on comparisons of pairwise correlations with DMS scores, see ‘Methods’. Error bars denote the 95% confidence interval of a binomial test.

Using either ΔΔG or |ΔΔG| values results in very similar rankings overall. In both cases, FoldX and Rosetta values derived on protein complexes were the top performers by a large margin, with FoldX monomer values ranking a distant third. Rankings starting at fifth place change slightly depending on whether we are using ΔΔG or | ΔΔG | values, but in both cases the four methods that show the least correspondence with DMS rankings are DynaMut2, mCSM, ENCoM for complexes and ENCoM for monomers.

Although a comprehensive benchmarking of VEPs using DMS datasets has been recently carried out^39^, we decided to compare how well predictors, which have been intentionally derived for functional variant effect evaluation, agree with experimental DMS scores on our structural benchmarking dataset. Many of our DMS dataset genes are not of human origin, and many VEPs are not designed to produce predictions for non-human proteins. We thus picked a small selection of methodologically diverse methods that had produced a sufficient number of predictions for most DMS datasets. This included two simple substitution matrices as baseline methods for assessing amino acid similarity, the widely used classical method SIFT^58^, more modern methods PROVEAN^59^ and SNAP2^60^, as well as state-of-the-art predictors DeepSequence^61^ and EVcouplings^62^. **Figure S1** shows that, compared to stability predictors, VEP correlations are considerably higher and more consistent across the different gene datasets. They achieve average Spearman’s correlations of 0.35, and 0.4 if we exclude the BLOSUM62 and Grantham substitution matrices^63,64^ – considerably higher than the average value of ~0.25 observed for stability predictors. If we include VEPs in our ranking scheme, we see that FoldX and Rosetta outperform the substitution matrices and SIFT, but cannot perform as well as modern VEPs for predicting functional scores from DMS experiments **(Figure S2),** consistent with our previous observation that stability predictors are, overall, less useful than VEPs for the identification of pathogenic missense mutations^12^.

### Computational stability predictions show better agreement with DMS scores derived through protein abundance-based assays

We have observed that the highest heterogeneity in correlation values arises not between different predictors, but across DMS datasets. We note that phenotypes that tend to correlate well with stability prediction values come from VAMP-seq or other fluorescence-based experiments (e.g., NUDT15, PTEN(a), TPMT, CYPC9(a), UBI4(b)). While growth assay datasets from some specific targets, like bla or P53, also show good agreement, they are uncharacteristic and more likely represent good protein-specific performance unrelated to the underlying assay type. Abundance-based approaches could be said to directly relate to the stability of target proteins, thus being better tailored to detecting loss of protein function. VAMP-seq has been specifically developed for this purpose and shown to correlate well with experimental thermodynamic and predicted stability measures^41,65^. On the other hand, competitive growth assays produce fitness values that relate to numerous underlying effects and various molecular mechanisms and show mixed tendencies for both high and low correlations, depending on the target.

To more quantitatively explore whether any specific assay types stand out in their agreement with stability prediction values, we classified the DMS datasets broadly into 5 groups, as outlined in **Supplementary Table 3,** depending on the phenotype assaying approach. Abundance-based assays represent experiments that make use of VAMP-seq and other fluorescent-tagging approaches that can quantify protein expression and stability in cells, being able to directly assess whether variants lead to a loss-of-function through degradation. Growth experiments involve mutant competition or antibiotic survival and are able to characterize functional effects from multiple molecular mechanisms. Binding assays such as phage display or two-hybrid experiments can represent variant impacts on intermolecular interactions, which ought to relate well to stability predictions for tools that can take into account relevant complex structures. The activity assay category includes target-specific experimental setups more directly measuring the functional mutant effects, and not just through the proxy of growth or protein stability. We separated out viral protein datasets, based on replication assays, into a separate category due to them being quite dissimilar from all other proteins.

**Figure 5** shows the mean per-dataset Spearman’s correlations with the top-ranking stability predictors FoldX and Rosetta, as well as the mean across all methods, for each assay phenotype group. The trend clearly demonstrates that stability predictors best reflect functional scores from abundance-based DMS datasets, both in the case of only the best methods, as well as for all predictors, while viral replication datasets show the worst agreement. Competitive growth assays, which are a popular generalized approach, appear to show mixed agreement on a per-gene level, but are not well correlated with changes to stability overall, possibly owing to destabilizing loss-of-function mechanisms being more prominent only in certain genes. Curiously, half of the binding-based assays do not show good agreement even with FoldX or Rosetta, which are able to evaluate interactions in complex structures, or show a marked improvement between predictions derived on monomeric structures *vs* fullest available structure.

**Figure 5.**
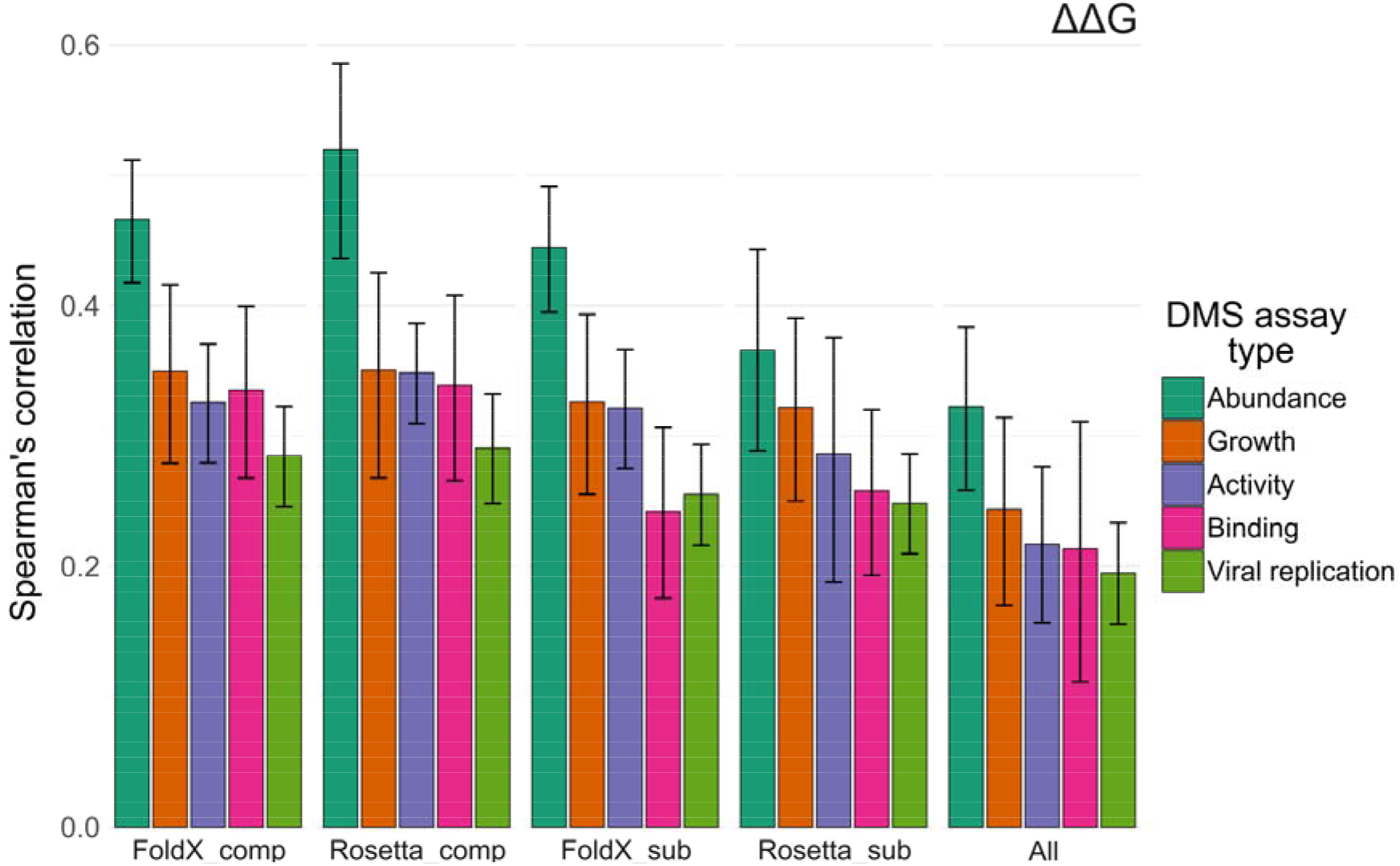
Stability predictions correlate best with results from assays interrogating antibiotic resistance and protein abundance. Correlations were calculated per-dataset, and the values shown represent assay group means. Error bars denote the 95% confidence interval, found by using the Fisher z transform of the correlation. Only raw ΔΔG values from stability predictors were used in the comparison.

*We* investigated the coverage of mapped and other available structures for the binding DMS proteins (PSD95, HRAS, GAL4, BRCA1), to see what factors might underly the low correlations. We found that structures for HRAS (PDB IDs: 2CE2 and 6POZ) and PSD95 (PDB ID: 6QJL) did not contain the binding partners relevant for the specific DMS assays, or were monomeric, explaining the inability of stability predictors to reflect intermolecular interactions. More suitable complex structures were not currently available in the PDB. However, both GAL4 and BRCA1(a) DMS datasets show increased correlation with stability values, when using complex structures. The GAL4 structure (PDB ID: 3COQ) contains a transcription factor dimer that is bound to a DNA duplex, allowing us to evaluate variant effects both on dimerization and DNA recognition. The BRCA1(a) dataset was derived from an assay interrogating BRCA1-BARD1 heterodimer interactions, which are represented in the structural data (PDB IDs: 7JZV and 1JM7).

Considering the strong correlation differences that may arise based on assay type, we decided to investigate how the ranking is affected by only using DMS datasets that have been shown to be well reflected by stability predictions. We explored DMS scores from experiments on protein abundance, which include VAMP-seq experiments and other fluorescence-based approaches. In our comparison we also included VEPs to see how stability predictors compare on DMS datasets that are best suited for loss-of-function prediction. **Figure 6** shows a considerably steeper rank distribution, strongly establishing Rosetta and FoldX predictions as the most reflective of functional assay scores between stability-based methods, with complex values for both leading the ranking. Importantly, complex-based Rosetta scores also outperform all the tested VEPs, although the slight advantage is statistically insignificant among the best methods. Overall, FoldX and Rosetta demonstrate similar performance on DMS experiments based on abundance phenotypes, and, judging by the overlapping confidence intervals, show a competitive performance with top VEPs on these datasets. While this current generation of stability predictors are not likely to be particularly useful in the direct identification of pathogenic variants compared to state-of-the-art VEPs, they offer a clear advantage over VEPs in terms of interpretability of mechanistic effects, especially in this case where we see that raw stability predictions correlate better than absolute values with functional DMS scores, indicating the distinction between stabilizing and destabilizing variants is important for accuracy. This contrasts with our past results for stability predictor performance in a classification task between pathogenic and benign variants, where |ΔΔG | values, focusing on the overall magnitude of a mutation’s impact on stability, showed better predictivity.

**Figure 6.**
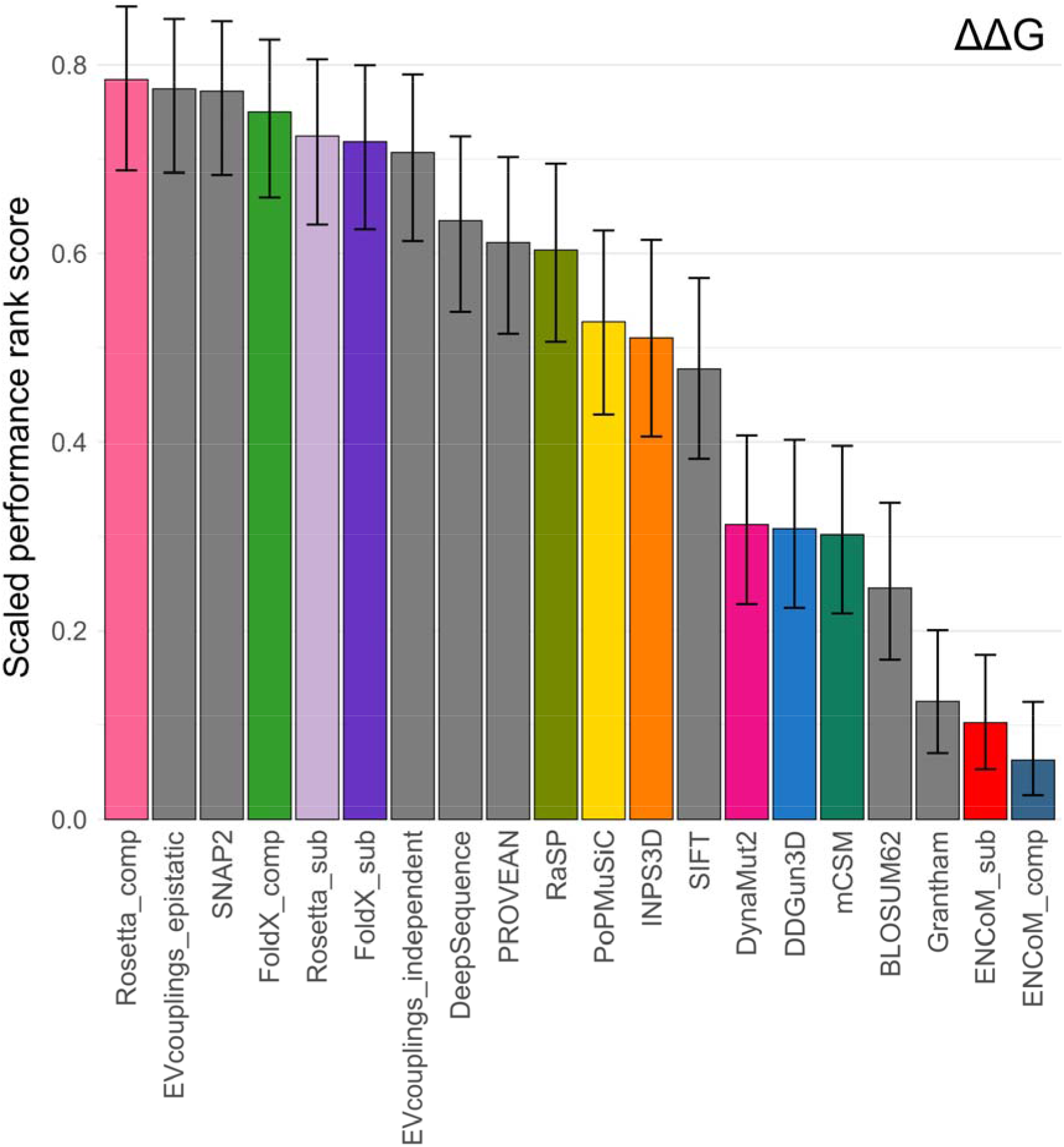
Computational stability predictor correlation becomes competitive with top VEPs when compared only against abundance-based DMS. Computational protein variant effect predictor rankings were derived based on comparisons of pairwise correlations with DMS scores, see ‘Methods’. Only raw ΔΔG values from stability predictors were used in the comparison. Error bars denote the 95% confidence interval of a binomial test.

## Discussion

Our study primarily aimed to examine the relationship between stability, structure and function, by interrogating the capacity of different protein stability predictors to score variants in line with experimentally determined functional impacts. We found that FoldX and Rosetta show the strongest correlations with DMS measurements, and also demonstrated the importance of protein complex structures for evaluating the functional impact of variants to the fullest extent. DMS experiments proved to be instrumental for this benchmarking task, as their results are currently the most direct representation of the functional protein variant landscape. Our results support the choice of Rosetta as a predictive tool, which has recently been used in similar studies exploring DMS functional score relationships with stability- and conservation-based metrics^48,50^. Unlike many VEPs, which are optimized for human mutations, or influenced by the widely varying sequence coverage across evolutionary space, stability predictors should be well suited to evaluate proteins in an organism-agnostic manner, as most are grounded in physics or approximate physical terms by proxy. Finally, we must accentuate that stability predictors are not designed for disease variant identification, and their training and design is aimed at reproducing realistic ΔΔG values that reflect experimental thermostability changes, and not rank functional impacts. Thus, our predictor ranking does not necessarily imply that FoldX or Rosetta are the most accurate tools for predicting either the ΔΔG magnitude, or the direction of stability perturbing effects. Despite this, stability predictors are still routinely used in clinical research for variant prioritization and mechanistic interpretation^29,30,32,66^, mostly because of their high interpretability due to their structure-based nature, and exploring the ability of ΔΔG to reflect functional impact could lead to a more effective application of these methodologies for such purposes.

Interestingly, we have previously also demonstrated FoldX was the best out of all tested stability predictors at distinguishing between confirmed human disease variants and putatively benign ones through both ΔΔG and | ΔΔG |, while Rosetta did not rank as high on that particular task^12^. One aspect of FoldX performance that made it useful for disease identification was its tendency to assign excessively large stability perturbation scores to disease variants, due to the clashes they caused within structures, very effectively separating the score densities between putatively benign mutations and truly deleterious ones. However, in this current work we demonstrate that FoldX and Rosetta also have the capacity to maintain an accurate relative ranking of functional variant effects, and not just produce outlier scores for disease mutations. This is potentially of great benefit, as it shows such predictors can delineate between hypomorphic and full loss-of-function effects for a given protein, which could be useful when interpreting and prioritizing variants or patient genotypes.

Some heterogeneity in predictor performance is not surprising. Previous efforts to benchmark their accuracy in reproducing realistic ΔΔG values have revealed highly varied performance, owing to the different methodological approaches and biases. A likely reason for the success of FoldX and Rosetta are their empirical scoring functions, containing energetic and statistical terms parametrized based on experimental data. It is unlikely that the source of high FoldX and Rosetta performance is overtraining on test data, as we are benchmarking a different kind of performance altogether. Instead, the high agreement is likely due to an underlying association of loss-of-function mechanisms with the protein and phenotype in question. RaSP^44^, a sequence-based method, benefits indirectly through being parametrized against Rosetta predictions, while leveraging considerably improved computational speed through a simplified model, beating other predictors that use sequence features. More novel approaches combining machine learning with sequence- or structure-based features, such as mCSM, DDGun3D, PoPMuSiC and INPS3D, do not seem to overall be as effective at ranking functional impacts. However, these state-of-the-art methods have been shown to be accurate at predicting actual ΔΔG values, and improve upon other methodologies in terms of prediction symmetry, and are less sensitive to resolution and substitution type biases^18,25–27,36^.

Interestingly, we observed the more unconventional methods, such as ENCoM and DynaMut2, benefit the most from using absolute ΔΔG values, but also, despite having adjusted for effect directions of predictors and DMS sets to match, often demonstrate moderate inverse correlations, likely due to experimental noise. It is also important to point that DDGun3D and INPS3D, the hybrid sequence and structure-based approaches, effectively include sequence-derived features representing evolutionary conservation in their predictions. Conservation is known to be predictive of damaging mutations, regardless of molecular mechanism, which would suggest that these methods could be capable of predicting mutations at conserved positions to be more (de)stabilizing than they actually are, resulting in a stronger correlation with the functional scores from DMS assays. Given that non-loss-of-function disease variants tend to be considerably structurally milder due to their molecular mechanisms, we would expect them to be poorly predicted by conventional stability predictors, since their damaging effects are unlikely to be due to destabilization^38^. DDGun3D and INPS3D appear to show relatively better correlations, compared with other methods, on essential, highly conserved genes, such as HSP82 and PAB1, or some genes with mixed disease mechanism like TARDBP and SRC^55–57^. However, sequence features do not appear to consistently favour the prediction of non-LOF disease gene variants, like CALM1, and overall these hybrid predictors demonstrate poor performance in the correlation ranking, and especially on the abundance-based DMS datasets.

In this study we also demonstrated the utility of using complex structures of biological units, as monomer structures may not be sufficient for assessing the full functional impact of a variant, for instance due to molecular mechanisms involving intermolecular interactions, as shown previously^38^. Both FoldX and Rosetta, the best ranked methods, and ENCoM, the poorest, were able to take into account structures containing more than one protein chain, and in the case of FoldX and Rosetta also other biomolecules. All methods saw increased correlations with DMS values for some datasets which are based on phenotypes that involved assessing binding, aggregation, or for genes with known involvement in functional interactions. The correlation between FoldX and the transcription factor GAL4 saw considerable benefit, because FoldX is able to evaluate the stability perturbations involving DNA. Of course, such an approach depends on having available structures, and being able to assess the relevance of a given assembly for assessing a particular phenotype. While DeepMind have now made monomeric AlphaFold2 model structures of the whole proteome available to all, the Protein Data Bank still remains the main source of functionally relevant protein complexes.

Our work hints at the pervasiveness of destabilizing loss-of-function mechanisms throughout the functional variant landscape, and the importance of choosing the most informative phenotypes for DMS experiments. Both stability predictors and VEPs we tested performed the best on experiments interrogating protein abundance and stability, such as various assays involving fluorescent reporters or VAMP-seq, and some antibiotic survival experiments. However, such DMS datasets are less likely to accurately reflect impacts of variants associated with other molecular mechanisms. In the case of proteins with multiple functions or disease mechanism, multiple different assays would be required to gleam the full scope of a variant. This could be alleviated by using competitive growth assays, which allow to capture the broadest selection of functional effects from variants, however, the delineation of what molecular mechanism might be responsible for the increase or reduction in fitness is lost. Other general issues with DMS data are that it can be noisy, restricted to a small set of experimental conditions, and also removed from the original cellular context^67^. Due to these limitations, as well as the enormous resource cost of most current DMS methodologies, they are unlikely to replace computational prediction tools as the main avenue to fully understanding functional effects of missense mutations in the near future. However, an exciting new methodology, dubbed cDNA display proteolysis, was recently shown to be capable of assessing functional variant effects on protein thermodynamic stability at tremendous scale and speed^68^. While limited to a stability phenotype, such a DMS approach also presents a valuable opportunity to gleam insight into the mechanisms of LOF disease, further test the accuracy of current computational tools on a large independent dataset and use it for training and developing better methodologies.

## Methods

### Structural DMS variant dataset collection and mapping

DMS datasets were gathered from experimental research publications, previous computational works that had systemized them, and additional new datasets were downloaded from the MaveDB^7^. Publication references and MaveDB accession codes are outlined in **Supplementary Table 1.**

Protein Data Bank structures were selected through a procedure published previously^38^, choosing the first biological assembly for each structure as representative of the biologically relevant quaternary structure. The mutation mapping pipeline has been previously described^38^. Protein chains with more than 90% sequence identity to a human protein over a region of at least 50 amino acid residues were considered. Mutations were only mapped to non-human structures in cases where the residue and its adjacent neighbours were the same as the human wild-type sequence. Including non-human structures with this approach allowed us to substantially increase the size of our dataset. Structures with best resolution followed by largest biological assembly were prioritized for mapping in the case of multiple available structures for a residue. For PDB files containing multiple occupancies of a single residue, only the first occurring entry was selected and residues missing from PDB structures were not considered. For NMR ensembles the first frame was chosen.

Variants that could not be mapped to PDB structures were evaluated on AlphaFold2 models^69^. AlphaFold models were accessed and downloaded on 2021.07.27 from https://alphafold.ebi.ac.uk. The DMS variants were mapped to AlphaFold models based on UniProt sequence positions.

### Structure-based variant stability predictors and variant effect predictors

FoldX 5.0 was run using default parameters as previously described^13,38^. Both protein subunit and complex structures were used where available to also take into account intermolecular interactions. The structures were passed through the ‘RepairPDB’ function prior to ΔΔG calculations. DDGun^16^ source code was downloaded from https://github.com/biofold/ddgun and run using the 3D protocol. ENCoM^15^ was run both on monomeric and complex PDB protein structures throughout two separate instances, source code was downloaded from https://github.com/NRGIab/ENCoM. The Cartesian ΔΔG application from Rosetta suite (Linux build 2021.16.61629) was run based on the protocol laid out in Park et al.^70^, while using the Ref2015 scoring function. The structures were relaxed according to the protocol for both monomeric and complex PDB structures. Results from three prediction iterations were averaged and ΔΔG values were derived by taking the difference between the wild-type and mutant values for each individual run. For complex evaluation, PDB structures 4JZW and 4JZZ were modified to introduce cysteines instead of some non-standard residues for Rosetta to recognize the disulfide bonds. RaSP^44^ was run on a modified Google Colab notebook, based on a mix of code from GitHub commits 26b0b1a and 518624e. INPS3D^47^, mCSM^17^, DynaMut2^46^ and PoPMuSiC^45^ webservers were queried in Python 3.8.8 using RoboBrowser (https://github.com/jmcarp/robobrowser) or Selenium (https://github.com/baijum/selenium-python) packages. Some predictions for certain datasets or variants could not be completed due to software or webserver errors.

VEP values were obtained using our previously described pipeline^39^. Where available, VEP predictions were obtained using the dbNSFP database version 4.0^71^. Further predictor scores were obtained from predictor web-interfaces like SNAP2^60^, or run locally for EVcouplings^62^, SIFT^58^ and DeepSequence^61^.

### Statistical analyses

Full-pairwise Spearman’s correlations for complete observations were calculated using the R ‘psych’^72^ and ‘corrplot’^73^ packages. The directions of the DMS scores and stability predictor values were adjusted to match a single ‘direction’.

For the predictor correlation ranking against DMS functional values, a scoring scheme involving pairwise comparisons of each predictor against every other predictor for every DMS dataset was derived. For a given DMS dataset and two predictors, only the common subset of variants, for which both prediction and assay values were available, was used to calculate Spearman’s rho values. The correlation values were rounded to two digits after the decimal point and were compared only if neither predictor was completely missing data for a given DMS dataset. After a successful comparison, the predictor with a larger correlation value was rewarded a point, or each predictor was granted half a point in the case of a draw after rounding. In the end, each predictor’s score was divided by the total number of successful comparisons it was involved in to normalize the scores. This normalization also gives a chance for predictors with a few missing DMS datasets not to fall behind in the scoring scheme due to undersampling issues. Error bars were derived using the ‘binom.confint’ function from the ‘binom’ package^74^.

## Data Availability

All the DMS dataset variants, structure identifiers used in the dataset, and stability predictor values are available at DOI: 10.17605/OSF.IO/ZTV8A

## Acknowledgements

This project was supported by the European Research Council (ERC) under the European Union’s Horizon 2020 research and innovation programme (grant agreement No. 101001169). JM is a Lister Institute Research Fellow.

## References

1. Dunham I, Kundaje A, Aldred SF, Collins PJ, Davis CA, Doyle F, Epstein CB, Frietze S, Harrow J, Kaul R, et al. (2012) An integrated encyclopedia of DNA elements in the human genome. Nature 489:57–74.

2. Karczewski KJ, Francioli LC, Tiao G, Cummings BB, Alföldi J, Wang Q, Collins RL, Laricchia KM, Ganna A, Birnbaum DP, et al. (2020) The mutational constraint spectrum quantified from variation in 141,456 humans. Nature 581:434–443.

3. Landrum MJ, Lee JM, Riley GR, Jang W, Rubinstein WS, Church DM, Maglott DR (2014) ClinVar: Public archive of relationships among sequence variation and human phenotype. Nucleic Acids Research 42:980–985.

4. Iversen ES Jr, Couch FJ, Goldgar DE, Tavtigian SV, Monteiro ANA (2011) A Computational Method to Classify Variants of Uncertain Significance Using Functional Assay Data with Application to BRCA1. Cancer Epidemiology, Biomarkers & Prevention 20:1078–1088.

5. Fowler DM, Fields S (2014) Deep mutational scanning: a new style of protein science. Nat Methods 11:801–807.

6. Starita LM, Ahituv N, Dunham MJ, Kitzman JO, Roth FP, Seelig G, Shendure J, Fowler DM (2017) Variant Interpretation: Functional Assays to the Rescue. The American Journal of Human Genetics 101:315–325.

7. Esposito D, Weile J, Shendure J, Starita LM, Papenfuss AT, Roth FP, Fowler DM, Rubin AF (2019) MaveDB: an open-source platform to distribute and interpret data from multiplexed assays of variant effect. Genome Biology 20:223.

8. Kuang D, Weile J, Kishore N, Nguyen M, Rubin AF, Fields S, Fowler DM, Roth FP (2021) MaveRegistry: a collaboration platform for multiplexed assays of variant effect. Bioinformatics 37:3382–3383.

9. AVE Alliance Founding Members (2021) The Atlas of Variant Effects (AVE) Alliance: understanding genetic variation at nucleotide resolution. Available from: https://doi.org/10.5281/zenodo.4989960

10. Livesey BJ, Marsh JA (2022) Interpreting protein variant effects with computational predictors and deep mutational scanning. Dis Model Mech 15.

11. Iqbal S, Li F, Akutsu T, Ascher DB, Webb GI, Song J (2021) Assessing the performance of computational predictors for estimating protein stability changes upon missense mutations. Briefings in Bioinformatics 22:bbab184.

12. Gerasimavicius L, Liu X, Marsh JA (2020) Identification of pathogenic missense mutations using protein stability predictors. Scientific Reports 10:1–10.

13. Delgado J, Radusky LG, Cianferoni D, Serrano L (2019) FoldX 5.0: Working with RNA, small molecules and a new graphical interface. Bioinformatics 35:4168–4169.

14. Alford RF, Leaver-Fay A, Jeliazkov JR, O’Meara MJ, DiMaio FP, Park H, Shapovalov MV, Renfrew PD, Mulligan VK, Kappel K, et al. (2017) The Rosetta All-Atom Energy Function for Macromolecular Modeling and Design. Journal of Chemical Theory and Computation 13:3031–3048.

15. Frappier V, Chartier M, Najmanovich RJ (2015) ENCoM server: Exploring protein conformational space and the effect of mutations on protein function and stability. Nucleic Acids Research 43:W395–W4OO.

16. Montanucci L, Capriotti E, Birolo G, Benevenuta S, Pancotti C, Lal D, Fariselli P (2022) DDGun: an untrained predictor of protein stability changes upon amino acid variants. Nucleic Acids Research 50:W222–W227.

17. Pires DEV, Ascher DB, Blundell TL (2014) MCSM: Predicting the effects of mutations in proteins using graph-based signatures. Bioinformatics 30:335–342.

18. Caldararu O, Blundell TL, Kepp KP (2021) A base measure of precision for protein stability predictors: structural sensitivity. BMC Bioinformatics 22:88.

19. Khan S, Vihinen M (2010) Performance of protein stability predictors. Human Mutation 31:675–684.

20. Lonquety M, Lacroix Z, Chomilier J (2007) BENCHMARKING STABILITY TOOLS: COMPARISON OF SOFTWARES DEVOTED TO PROTEIN STABILITY CHANGES INDUCED BY POINT MUTATIONS PREDICTION. Comput Sys Bioinf Conference CSB2007 San Diego, USA 1.

21. Marabotti A, Del Prete E, Scafuri B, Facchiano A (2021) Performance of Web tools for predicting changes in protein stability caused by mutations. BMC Bioinformatics 22:345.

22. Pancotti C, Benevenuta S, Birolo G, Alberini V, Repetto V, Sanavia T, Capriotti E, Fariselli P (2022) Predicting protein stability changes upon single-point mutation: a thorough comparison of the available tools on a new dataset. Briefings in Bioinformatics 23:bbab555.

23. Potapov V, Cohen M, Schreiber G (2009) Assessing computational methods for predicting protein stability upon mutation: Good on average but not in the details. Protein Engineering, Design and Selection 22:553–560.

24. König E, Rainer J, Domingues FS (2016) Computational assessment of feature combinations for pathogenic variant prediction. Molecular Genetics & Genomic Medicine 4:431–446.

25. Montanucci L, Savojardo C, Martelli PL, Casadio R, Fariselli P (2019) On the biases in predictions of protein stability changes upon variations: the INPS test case Valencia A, editor. Bioinformatics 35:2525–2527.

26. Pucci F, Bernaerts K V., Kwasigroch JM, Rooman M (2018) Quantification of biases in predictions of protein stability changes upon mutations. Bioinformatics (Oxford, England) 34:3659–3665.

27. Usmanova DR, Bogatyreva NS, Bernad JA, Eremina AA, Gorshkova AA, Kanevskiy GM, Lonishin LR, Meister AV, Yakupova AG, Kondrashov FA, et al. (2018) Self-consistency test reveals systematic bias in programs for prediction change of stability upon mutation. Bioinformatics 34:3653–3658.

28. Blanco JD, Radusky L, Climente-González H, Serrano L (2018) FoldX accurate structural protein-DNA binding prediction using PADA1 (Protein Assisted DNA Assembly 1). Nucleic Acids Research 46:3852–3863.

29. Heyn P, Logan CV, Fluteau A, Challis RC, Auchynnikava T, Martin CA, Marsh JA, Taglini F, Kilanowski F, Parry DA, et al. (2019) Gain-of-function DNMT3A mutations cause microcephalic dwarfism and hypermethylation of Polycomb-regulated regions. Nature Genetics 51:96–105.

30. Holt RJ, Young RM, Crespo B, Ceroni F, Curry CJ, Bellacchio E, Bax DA, Ciolfi A, Simon M, Fagerberg CR, et al. (2019) De Novo Missense Variants in FBXW11 Cause Diverse Developmental Phenotypes Including Brain, Eye, and Digit Anomalies. American Journal of Human Genetics 105:640–657.

31. Othman H, Bouslama Z, Brandenburg J-T, da Rocha J, Hamdi Y, Ghedira K, Srairi-Abid N, Hazelhurst S (2020) Interaction of the spike protein RBD from SARS-CoV-2 with ACE2: Similarity with SARS-CoV, hot-spot analysis and effect of the receptor polymorphism. Biochemical and Biophysical Research Communications 527:702–708.

32. Williamson KA, Hall HN, Owen LJ, Livesey BJ, Hanson IM, Adams GGW, Bodek S, Calvas P, Castle B, Clarke M, et al. (2020) Recurrent heterozygous PAX6 missense variants cause severe bilateral microphthalmia via predictable effects on DNA–protein interaction. Genetics in Medicine 22:598–609.

33. Zhao Y, Li D, Bai X, Luo M, Feng Y, Zhao Y, Ma F, Yang G-Y (2021) Improved thermostability of proteinase K and recognizing the synergistic effect of Rosetta and FoldX approaches. Protein Engineering, Design and Selection 34:gzab024.

34. Fu H, Siggs O, Knight L, Staffieri SE, Ruddle JB, Birsner AE, Collantes ER, Craig J, Wiggs JL, D’Amato RJ (2022) Thrombospondin-1 p.R1034 missense alleles cause congenital glaucoma with variable expressivity by inducing extracellular protein aggregation. Investigative Ophthalmology & Visual Science 63:805–805.

35. Yu Z, Yu H, Xu J, Wang Z, Wang Z, Kang T, Chen K, Pu Z, Wu J, Yang L, et al. (2022) Enhancing thermostability of lipase from Pseudomonas alcaligenes for producing l-menthol by the CREATE strategy. Catal. Sci. Technol. 12:2531–2541.

36. Sanavia T, Birolo G, Montanucci L, Turina P, Capriotti E, Fariselli P (2020) Limitations and challenges in protein stability prediction upon genome variations: towards future applications in precision medicine. Computational and Structural Biotechnology Journal 18:1968–1979.

37. Birolo G, Benevenuta S, Fariselli P, Capriotti E, Giorgio E, Sanavia T (2021) Protein Stability Perturbation Contributes to the Loss of Function in Haploinsufficient Genes. Front. Mol. Biosci. 8:620793.

38. Gerasimavicius L, Livesey BJ, Marsh JA (2022) Loss-of-function, gain-of-function and dominant negative mutations have profoundly different effects on protein structure. Nature Communications 13:3895.

39. Livesey BJ, Marsh JA (2020) Using deep mutational scanning to benchmark variant effect predictors and identify disease mutations. Molecular Systems Biology 16:1–12.

40. Livesey BJ, Marsh JA (2022) Updated benchmarking of variant effect predictors using deep mutational scanning. bioRxiv:2022.11.19.517196.

41. Matreyek KA, Starita LM, Stephany JJ, Martin B, Chiasson MA, Gray VE, Kircher M, Khechaduri A, Dines JN, Hause RJ, et al. (2018) Multiplex assessment of protein variant abundance by massively parallel sequencing. Nat Genet 50:874–882.

42. Zheng H, Yan X, Li G, Lin H, Deng S, Zhuang W, Yao F, Lu Y, Xia X, Yuan H, et al. (2022) Proactive functional classification of all possible missense single-nucleotide variants in KCNQ4. Genome Res.

43. Akdel M, Pires DEV, Porta Pardo E, Jänes J, Zalevsky AO, Mészáros B, Bryant P, Good LL, Laskowski RA, Pozzati G, et al. (2021) A structural biology community assessment of AlphaFold 2 applications. bioRxiv:2021.09.26.461876.

44. Blaabjerg LM, Kassem MM, Good LL, Jonsson N, Cagiada M, Johansson KE, Boomsma W, Stein A, Lindorff-Larsen K (2022) Rapid protein stability prediction using deep learning representations. bioRxiv:2022.07.14.500157.

45. Dehouck Y, Kwasigroch JM, Gilis D, Rooman M (2011) PoPMuSiC 2.1: A web server for the estimation of protein stability changes upon mutation and sequence optimality. BMC Bioinformatics 12:151.

46. Rodrigues CHM, Pires DEV, Ascher DB (2021) DynaMut2: Assessing changes in stability and flexibility upon single and multiple point missense mutations. Protein Science 30:60–69.

47. Savojardo C, Fariselli P, Martelli PL, Casadio R (2016) INPS-MD: a web server to predict stability of protein variants from sequence and structure. Bioinformatics 32:2542–2544.

48. Høie MH, Cagiada M, Beck Frederiksen AH, Stein A, Lindorff-Larsen K (2022) Predicting and interpreting large-scale mutagenesis data using analyses of protein stability and conservation. Cell Reports 38:110207.

49. Montanucci L, Martelli PL, Ben-Tal N, Fariselli P (2019) A natural upper bound to the accuracy of predicting protein stability changes upon mutations Valencia A, editor. Bioinformatics 35:1513–1517.

50. Cagiada M, Johansson KE, Valanciute A, Nielsen SV, Hartmann-Petersen R, Yang JJ, Fowler DM, Stein A, Lindorff-Larsen K (2021) Understanding the Origins of Loss of Protein Function by Analyzing the Effects of Thousands of Variants on Activity and Abundance Ozkan B, editor. Molecular Biology and Evolution 38:3235–3246.

51. Rocchetti M, Sala L, Dreizehnter L, Crotti L, Sinnecker D, Mura M, Pane LS, Altomare C, Torre E, Mostacciuolo G, et al. (2017) Elucidating arrhythmogenic mechanisms of long-QT syndrome CALM1-F142L mutation in patient-specific induced pluripotent stem cell-derived cardiomyocytes. Cardiovascular Research 113:531–541.

52. Badone B, Ronchi C, Kotta M-C, Sala L, Ghidoni A, Crotti L, Zaza A (2018) Calmodulinopathy: Functional Effects of CALM Mutations and Their Relationship With Clinical Phenotypes. Frontiers in Cardiovascular Medicine [Internet] 5. Available from: https://www.frontiersin.org/articles/10.3389/fcvm.2018.00176

53. Boczek NJ, Gomez-Hurtado N, Ye D, Calvert ML, Tester DJ, Kryshtal DO, Hwang HS, Johnson CN, Chazin WJ, Loporcaro CG, et al. (2016) Spectrum and Prevalence of CALM1-, CALM2-, and CALM3-Encoded Calmodulin Variants in Long QT Syndrome and Functional Characterization of a Novel Long QT Syndrome-Associated Calmodulin Missense Variant, E141G. Circulation: Cardiovascular Genetics 9:136–146.

54. Suk TR, Rousseaux MWC (2020) The role of TDP-43 mislocalization in amyotrophic lateral sclerosis. Molecular Neurodegeneration 15:45.

55. Kabashi E, Lin L, Tradewell ML, Dion PA, Bercier V, Bourgouin P, Rochefort D, Bel Hadj S, Durham HD, Velde CV, et al. (2010) Gain and loss of function of ALS-related mutations of TARDBP (TDP-43) cause motor deficits in vivo. Human Molecular Genetics 19:671–683.

56. Turro E, Greene D, Wijgaerts A, Thys C, Lentaigne C, Bariana TK, Westbury SK, Kelly AM, Selleslag D, Stephens JC, et al. (2016) A dominant gain-of-function mutation in universal tyrosine kinase SRC causes thrombocytopenia, myelofibrosis, bleeding, and bone pathologies. Science Translational Medicine 8:328ra30–328ra30.

57. Abe K, Cox A, Takamatsu N, Velez G, Laxer RM, Tse SML, Mahajan VB, Bassuk AG, Fuchs H, Ferguson PJ, et al. (2019) Gain-of-function mutations in a member of the Src family kinases cause autoinflammatory bone disease in mice and humans. Proceedings of the National Academy of Sciences 116:11872–11877.

58. Ng PC, Henikoff S (2003) SIFT: predicting amino acid changes that affect protein function. Nucleic Acids Res 31:3812–3814.

59. Choi Y, Chan AP (2015) PROVEAN web server: A tool to predict the functional effect of amino acid substitutions and indels. Bioinformatics 31:2745–2747.

60. Hecht M, Bromberg Y, Rost B (2015) Better prediction of functional effects for sequence variants. BMC Genomics 16.

61. Riesselman AJ, Ingraham JB, Marks DS (2018) Deep generative models of genetic variation capture the effects of mutations. Nature Methods 15:816–822.

62. Hopf TA, Green AG, Schubert B, Mersmann S, Schärfe CPI, Ingraham JB, Toth-Petroczy A, Brock K, Riesselman AJ, Palmedo P, et al. (2019) The EVcouplings Python framework for coevolutionary sequence analysis. Bioinformatics 35:1582–1584.

63. Henikoff S, Henikoff JG (2000) Amino acid substitution matrices. Advances in Protein Chemistry 54:73–97.

64. Grantham R (1974) Amino acid difference formula to help explain protein evolution. Science 185:862–864.

65. Suiter CC, Moriyama T, Matreyek KA, Yang W, Scaletti ER, Nishii R, Yang W, Hoshitsuki K, Singh M, Trehan A, et al. (2020) Massively parallel variant characterization identifies NUDT15 alleles associated with thiopurine toxicity. Proceedings of the National Academy of Sciences 117:5394–5401.

66. McEntagart M, Williamson KA, Rainger JK, Wheeler A, Seawright A, De Baere E, Verdin H, Bergendahl LT, Quigley A, Rainger J, et al. (2016) A Restricted Repertoire of de Novo Mutations in ITPR1 Cause Gillespie Syndrome with Evidence for Dominant-Negative Effect. American Journal of Human Genetics 98:981–992.

67. Chiasson M, Dunham MJ, Rettie AE, Fowler DM (2019) Applying Multiplex Assays to Understand Variation in Pharmacogenes. Clin Pharmacol Ther 106:290–294.

68. Tsuboyama K, Dauparas J, Chen J, Mangan NM, Ovchinnikov S, Rocklin GJ (2022) Mega-scale experimental analysis of protein folding stability in biology and protein design. bioRxiv:2022.12.06.519132.

69. Jumper J, Evans R, Pritzel A, Green T, Figurnov M, Ronneberger O, Tunyasuvunakool K, Bates R, Žídek A, Potapenko A, et al. (2021) Highly accurate protein structure prediction with AlphaFold. Nature 596:583–589.

70. Park H, Bradley P, Greisen P, Liu Y, Mulligan VK, Kim DE, Baker D, Dimaio F (2016) Simultaneous Optimization of Biomolecular Energy Functions on Features from Small Molecules and Macromolecules. Journal of Chemical Theory and Computation 12:6201–6212.

71. Liu X, Li C, Mou C, Dong Y, Tu Y (2020) dbNSFP v4: a comprehensive database of transcript-specific functional predictions and annotations for human nonsynonymous and splice-site SNVs. Genome Med 12:103.

72. Revelle W (2022) psych: Procedures for Psychological, Psychometric, and Personality Research. Available from: https://CRAN.R-project.org/package=psych

73. Wei T, Simko V, Levy M, Xie Y, Jin Y, Zemla J, Freidank M, Cai J, Protivinsky T (2021) corrplot: Visualization of a Correlation Matrix. Available from: https://CRAN.R-project.org/package=corrplot

74. Dorai-Raj S (2014) binom: Binomial Confidence Intervals For Several Parameterizations. Available from: https://CRAN.R-project.org/package=binom

