## Supplemental Material for "Correspondence between functional scores from deep mutational scans and predicted effects on protein stability"

Supplementary figures and tables

Lukas Gerasimavicius, Benjamin J Livesey and Joseph A. Marsh*

MRC Human Genetics Unit, Institute of Genetics & Cancer, University of Edinburgh, Edinburgh, UK

*

***Supplementary***
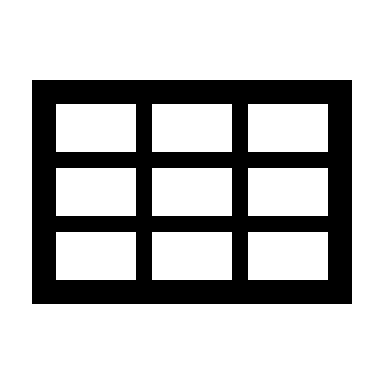
***Table 1. DMS datasets used in this study.***

| Uniprot ID | Dataset | Organism | Functional assay | Reference* |
| --- | --- | --- | --- | --- |
| P07550 | ADRB2 | Human | Pathway-specific reporter RNA-seq | ^1^ |
| P62593 | bla_a | E. coli | Antibiotic resistance assay | 00000070-a-2 |
| P62593 | bla_b | E. coli | Antibiotic resistance assay | 00000086-e-1 |
| P62593 | bla_c | E. coli | Antibiotic resistance assay | ^2^ |
| P62593 | bla_d | E. coli | Statistical ΔΔG from antibiotic resistance assay | ^3^ |
| P38398 | BRCA1_a | Human | Phage display | ^4^ |
| P38398 | BRCA1_b | Human | Competitive growth in human cells | ^5^ |
| P0DP23 | CALM1 | Human | Competitive growth in yeast | ^6^ |
| Q99ZW2 | Cas9 | S. pyrogenes | Survival assay | ^7^ |
| P62554 | ccdB | E. coli | Toxicity assay | ^8^ |
| P11712 | CYP2C9_a | Human | Clic-seq in humanized yeast | 00000095-a-1 |
| P11712 | CYP2C9_b | Human | Vamp-seq | 00000095-b-1 |
| P03377 | env | HIV virus | Competitive replication assay | ^9^ |
| P04386 | GAL4 | Yeast | Two-hybrid assay | ^10^ |
| P35557 | GCK | Human | Functional complementation in yeast | 00000096-a-1 |
| P31150 | GDI1 | Human | Functional complementation in yeast | 00000066-a-1 |
| Custom seq. | GmR | E. coli | Antibiotic resistance | ^11^ |
| P20589 | haeIIIM | H. aegyptius | Competitive growth | ^12^ |
| A0A2Z5U3Z0 | H1N1 | Influenza virus | Competitive replication assay | ^13^ |
| A0A097PF60 | H3N2 | Influenza virus | Competitive replication assay | ^14^ |
| P01112 | HRAS | Human | Two-hybrid assay | ^15^ |
| P02829 | HSP82 | Yeast | Competitive growth assay | ^16^ |
| P69222 | infA | E. coli | Competitive growth assay | ^17^ |
| P56696 | KCNQ4 | Human | Patch-clamp whole-cell current assay | 00000094-a-2 |
| P28482 | MAPK1 | Human | Competitive growth assay | ^18^ |
| P43246 | MSH2 | Human | MMR activity in human HAP1 cells | 00000050-a-1 |
| Q9NV35 | NUDT15 | Human | Meta-analysis of VAMP-seq and thiopurine-cytotoxicity assays | 00000055-0-1 |
| P04637 | P53_a | Human | Competitive growth assay | ^19^ |
| P04637 | P53_b | Human | 3D culture relative proliferation assay in human cells | 00000059-a-1 |
| P15659 | PA | Influenza virus | Competitive replication assay | ^20^ |
| P04147 | PAB1 | Yeast | Competitive growth | ^21^ |
| P78352 | PSD95 | Human | Bacterial two-hybrid assay | 00000053-a-2 |
| P60484 | PTEN_a | Human | Vamp-seq | 00000013-a-1 |
| P60484 | PTEN_b | Human | Lipid phosphatase assay in humanized yeast | 00000054-a-1 |
| P12931 | SRC | Human | Growth assay in yeast | 00000041-a-1 |
| P63165 | SUMO1 | Human | Competitive growth in yeast | ^6^ |
| Q13148 | TARDBP_a | Human | Yeast toxicity assay | 00000060-a-1 |
| Q13148 | TARDBP_b | Human | Yeast toxicity assay | 00000060-a-2 |
| Q9H3S3 | TPK1 | Human | Competitive growth in yeast | ^6^ |
| P51580 | TPMT | Human | Vamp-seq | ^22^ |
| P63279 | UBE2I | Human | Competitive growth in yeast | ^6^ |
| P0CG63 | UBI4_a | Yeast | Competitive growth | ^23^ |
| P0CG63 | UBI4_b | Yeast | Fluorescence-based assay | ^24^ |
| Q9BQB6 | VKOR_a | Human | Carboxylation activity | 00000078-a-1 |
| Q9BQB6 | VKOR_b | Human | Vamp-seq | 00000078-b-1 |

*Citation or MaveDB ID

***Supplementary Table 2. The variant and prediction dataset used in this study, available as a file (Supplementary_Table_2.csv).***

Supplementary
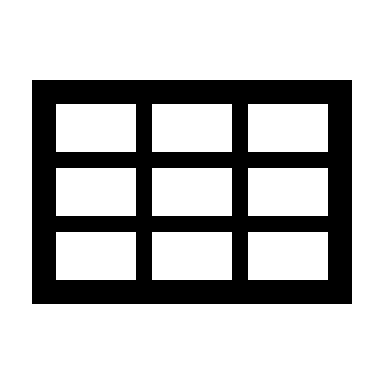
Table 3. DMS dataset classification by assay type.

| Abundance | CYP2C9_b, NUDT15, PTEN_a, TPMT, UBI4_b, VKOR_b |
| --- | --- |
| Activity | ADRB2, CYP2C9_a, GCK, KCNQ4, MSH2, PTEN_b, VKOR_a |
| Binding | BRCA1_a, GAL4, HRAS, PSD95 |
| Viral replication | env, H1N1, H3N2, PA |
| Growth | bla_a, bla_b, bla_c, bla_d, BRCA1_b, CALM1, Cas9, ccdB, GDI1, GmR, haeIIIM, HSP82, infA, MAPK1, P53_a, P53_b, PAB1, SRC, SUMO1, TARDBP_a, TARDBP_b, TPK1, UBE2I, UBI4_a |

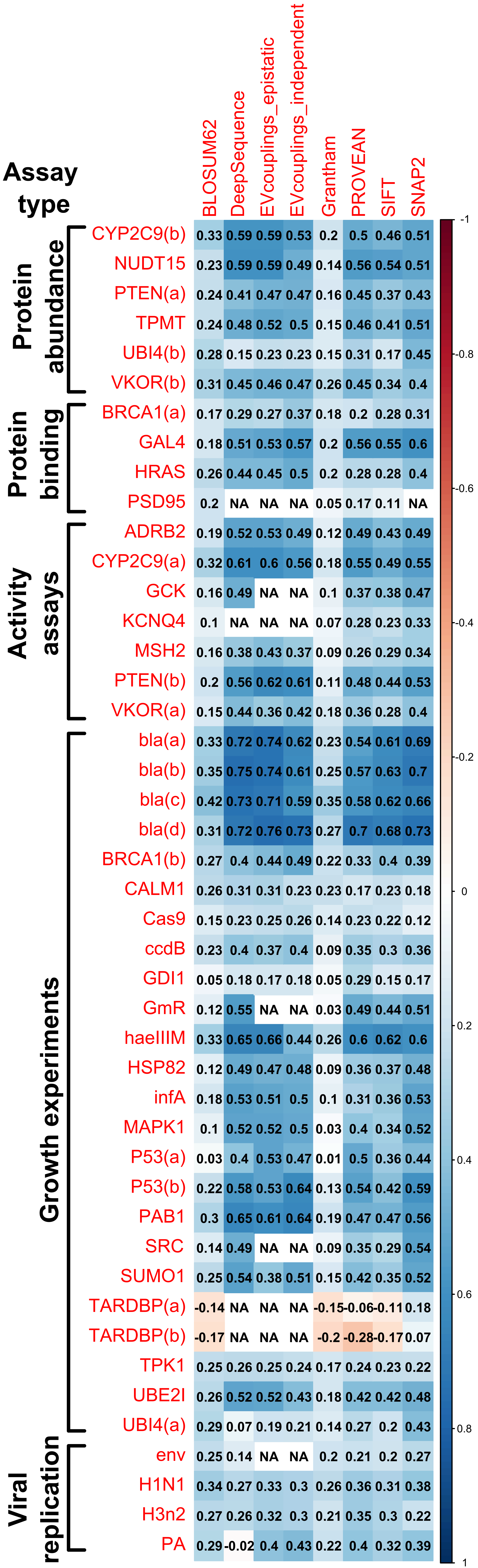

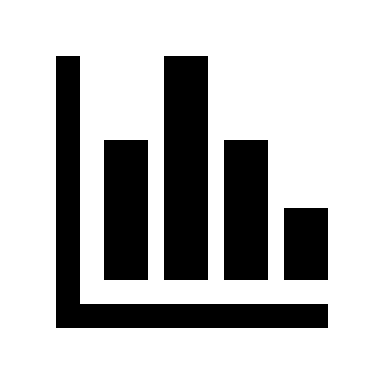
Figure S1. VEPs demonstrate a considerably higher degree of correlation to DMS values and performance homogeneity per-dataset than stability predictors. The coloured scale bar represents the absolute magnitude of Spearman’s rho values from pairwise comparisons calculated for complete observations. All predictor ΔΔG values were adjusted to match for stabilizing and destabilizing effect directions.

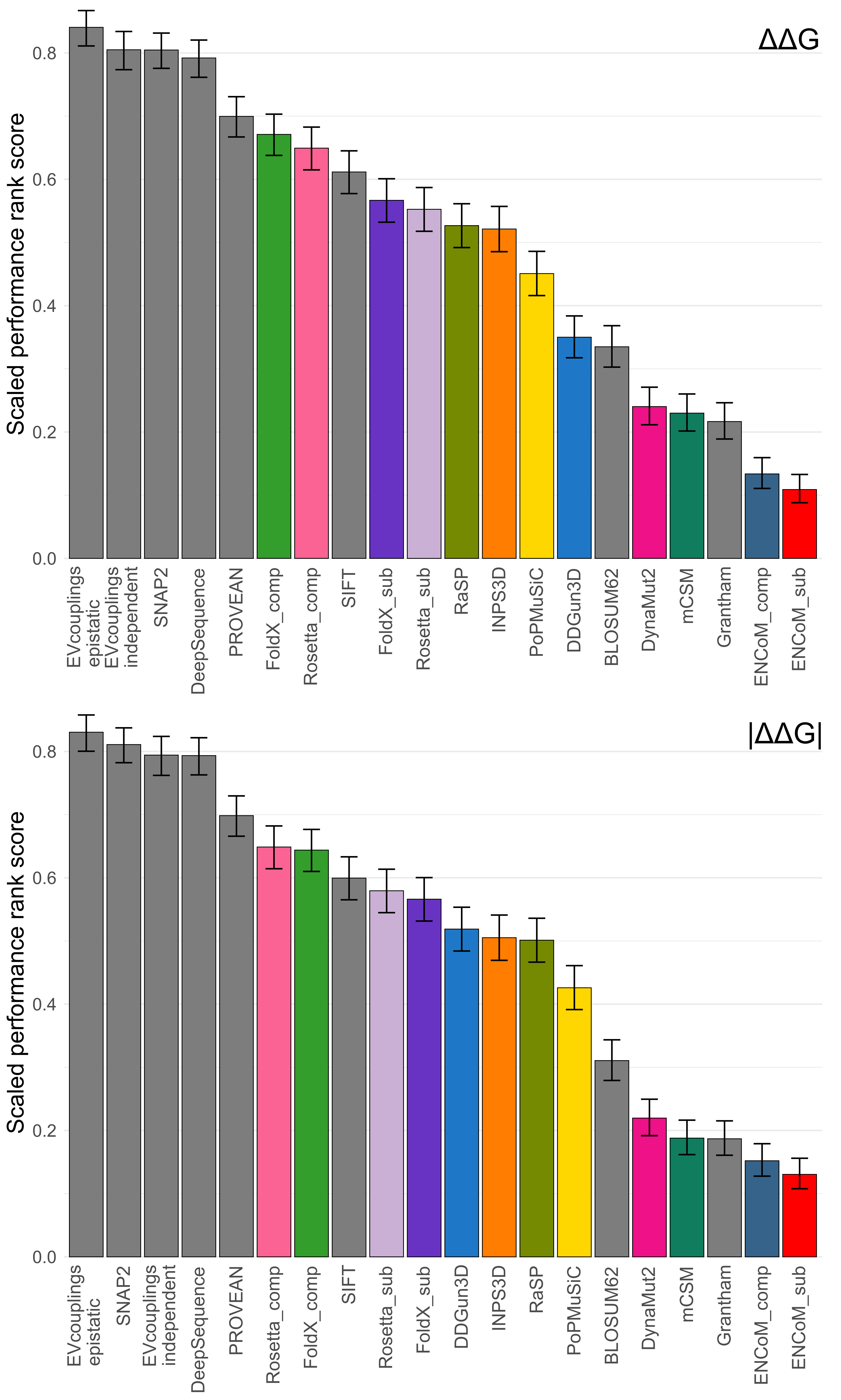

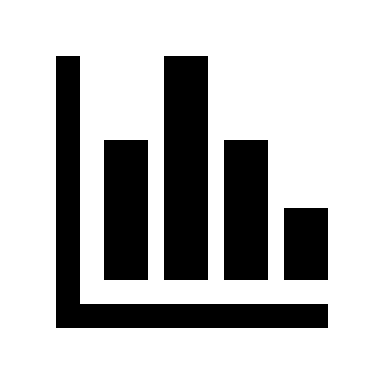
Figure S2. Most tested VEPs outperform protein stability predictors in ranking variant effects based on DMS scores. Computational protein variant effect predictor rankings were derived based on comparisons of pairwise correlations with DMS scores see ‘Methods’. Error bars denote the 95% confidence interval of a binomial test.

### References

1. Jones EM, Lubock NB, Venkatakrishnan A, Wang J, Tseng AM, Paggi JM, Latorraca NR, Cancilla D, Satyadi M, Davis JE, et al. (2020) Structural and functional characterization of G protein–coupled receptors with deep mutational scanning Larhammar D, Aldrich RW, Fraser JS, Manglik A, editors. eLife 9:e54895.

2. Jacquier H, Birgy A, Le Nagard H, Mechulam Y, Schmitt E, Glodt J, Bercot B, Petit E, Poulain J, Barnaud G, et al. (2013) Capturing the mutational landscape of the beta-lactamase TEM-1. Proceedings of the National Academy of Sciences 110:13067–13072.

3. Deng Z, Huang W, Bakkalbasi E, Brown NG, Adamski CJ, Rice K, Muzny D, Gibbs RA, Palzkill T (2012) Deep Sequencing of Systematic Combinatorial Libraries Reveals β-Lactamase Sequence Constraints at High Resolution. Journal of Molecular Biology 424:150–167.

4. Starita LM, Young DL, Islam M, Kitzman JO, Gullingsrud J, Hause RJ, Fowler DM, Parvin JD, Shendure J, Fields S (2015) Massively Parallel Functional Analysis of BRCA1 RING Domain Variants. Genetics 200:413–422.

5. Findlay GM, Daza RM, Martin B, Zhang MD, Leith AP, Gasperini M, Janizek JD, Huang X, Starita LM, Shendure J (2018) Accurate classification of BRCA1 variants with saturation genome editing. Nature 562:217–222.

6. Weile J, Sun S, Cote AG, Knapp J, Verby M, Mellor JC, Wu Y, Pons C, Wong C, Lieshout N, et al. (2017) A framework for exhaustively mapping functional missense variants. Molecular Systems Biology 13:957.

7. Spencer JM, Zhang X (2017) Deep mutational scanning of S. pyogenes Cas9 reveals important functional domains. Scientific Reports 7:16836.

8. Adkar BV, Tripathi A, Sahoo A, Bajaj K, Goswami D, Chakrabarti P, Swarnkar MK, Gokhale RS, Varadarajan R (2012) Protein Model Discrimination Using Mutational Sensitivity Derived from Deep Sequencing. Structure 20:371–381.

9. Haddox HK, Dingens AS, Bloom JD (2016) Experimental Estimation of the Effects of All Amino-Acid Mutations to HIV’s Envelope Protein on Viral Replication in Cell Culture. PLOS Pathogens 12:e1006114.

10. Kitzman JO, Starita LM, Lo RS, Fields S, Shendure J (2015) Massively parallel single-amino-acid mutagenesis. Nature Methods 12:203–206.

11. Dandage R, Pandey R, Jayaraj G, Rai M, Berger D, Chakraborty K (2018) Differential strengths of molecular determinants guide environment specific mutational fates. PLOS Genetics 14:e1007419.

12. Rockah-Shmuel L, Tóth-Petróczy Á, Tawfik DS (2015) Systematic Mapping of Protein Mutational Space by Prolonged Drift Reveals the Deleterious Effects of Seemingly Neutral Mutations. PLOS Computational Biology 11:e1004421.

13. Doud MB, Bloom JD (2016) Accurate Measurement of the Effects of All Amino-Acid Mutations on Influenza Hemagglutinin. Viruses 8.

14. Lee JM, Huddleston J, Doud MB, Hooper KA, Wu NC, Bedford T, Bloom JD (2018) Deep mutational scanning of hemagglutinin helps predict evolutionary fates of human H3N2 influenza variants. Proceedings of the National Academy of Sciences 115:E8276–E8285.

15. Bandaru P, Shah NH, Bhattacharyya M, Barton JP, Kondo Y, Cofsky JC, Gee CL, Chakraborty AK, Kortemme T, Ranganathan R, et al. (2017) Deconstruction of the Ras switching cycle through saturation mutagenesis Valencia A, editor. eLife 6:e27810.

16. Mishra P, Flynn JM, Starr TN, Bolon DNA (2016) Systematic Mutant Analyses Elucidate General and Client-Specific Aspects of Hsp90 Function. Cell Reports 15:588–598.

17. Kelsic ED, Chung H, Cohen N, Park J, Wang HH, Kishony R (2016) RNA Structural Determinants of Optimal Codons Revealed by MAGE-Seq. Cell Systems 3:563-571.e6.

18. Brenan L, Andreev A, Cohen O, Pantel S, Kamburov A, Cacchiarelli D, Persky NS, Zhu C, Bagul M, Goetz EM, et al. (2016) Phenotypic Characterization of a Comprehensive Set of MAPK1/ERK2 Missense Mutants. Cell Reports 17:1171–1183.

19. Giacomelli AO, Yang X, Lintner RE, McFarland JM, Duby M, Kim J, Howard TP, Takeda DY, Ly SH, Kim E, et al. (2018) Mutational processes shape the landscape of TP53 mutations in human cancer. Nature Genetics 50:1381–1387.

20. Wu NC, Olson CA, Du Y, Le S, Tran K, Remenyi R, Gong D, Al-Mawsawi LQ, Qi H, Wu T-T, et al. (2015) Functional Constraint Profiling of a Viral Protein Reveals Discordance of Evolutionary Conservation and Functionality. PLOS Genetics 11:e1005310.

21. Melamed D, Young DL, Gamble CE, Miller CR, Fields S (2013) Deep mutational scanning of an RRM domain of the Saccharomyces cerevisiae poly(A)-binding protein. RNA 19:1537–1551.

22. Matreyek KA, Starita LM, Stephany JJ, Martin B, Chiasson MA, Gray VE, Kircher M, Khechaduri A, Dines JN, Hause RJ, et al. (2018) Multiplex assessment of protein variant abundance by massively parallel sequencing. Nat Genet 50:874–882.

23. Roscoe BP, Thayer KM, Zeldovich KB, Fushman D, Bolon DNA (2013) Analyses of the Effects of All Ubiquitin Point Mutants on Yeast Growth Rate. Journal of Molecular Biology 425:1363–1377.

24. Roscoe BP, Bolon DNA (2014) Systematic Exploration of Ubiquitin Sequence, E1 Activation Efficiency, and Experimental Fitness in Yeast. Journal of Molecular Biology 426:2854–2870.
